# Interdependency between oxytocin and dopamine in trust-based learning in mice

**DOI:** 10.1101/2025.06.11.659016

**Authors:** Samuel Budniok, Zsuzsanna Callaerts-Vegh, Marian Bakermans-Kranenburg, Guy Bosmans, Rudi D’Hooge

**Author notes:** These authors contributed equally to this work and share last authorship (Guy Bosmans, Rudi D’Hooge). Corresponding author: Samuel Budniok Laboratory of Biological Psychology Faculty of Psychology and Educational Sciences University of Leuven Tiensestraat 102, 3000 Leuven, Belgium.

## Abstract

Oxytocin (OT) is a neuropeptide implicated in complex social behaviors including trust and attachment, yet the neural mechanisms underlying its effects remain unclear. OT is thought to modulate behavior by enhancing the salience of social cues and attenuating prediction error (PE) processing, the discrepancy between expected and actual outcomes that drives learning. Since both salience coding and PE processing involve dopamine (DA) neurons as well, the current study investigated the putative interdependence between OT and DA in social safety learning using the social transmission of food preference (STFP) paradigm. STFP is based on the observation that mice (observers) display neophobia toward novel food, but develop a preference for it after a conspecific demonstrator signals its safety. We interpreted STFP acquisition as a functional parallel to human trust-based learning and found that OT enhanced learning in a trust acquisition condition, but only when DA signaling was intact. In a trust violation condition, where demonstrated food was later paired with lithium chloride (LiCl)-induced food aversion, both OT and DA depletion blocked learning, resulting in retained preference for demonstrated food, but not when OT was administered under DA depletion. These findings reveal a functional interaction between the OT and DA systems to modulate social safety learning, which may have important implications for OT’s potential in treating disorders involving DA dysfunction.

## Introduction

The hypothalamic neuropeptide oxytocin (OT), identified over a century ago by Sir Henry Dale [1], was initially defined by its hormonal effects on the reproductive system [2]. More recent research has implicated centrally released OT in the modulation of various social behaviors and socio-cognitive processes such as social recognition, trust and attachment [3–5]. Putative mechanisms through which OT exerts effects include enhancing the salience of social cues [6] and attenuating prediction error (PE) processing [7]. PE refers to the discrepancy between expected and actual outcomes that drives learning [8]. Notably, both salience coding and PE processing are thought to involve mesocorticolimbic dopamine (DA) neurons as well [9–10]. These neurons originate in the ventral tegmental area (VTA) and project broadly to cortical regions, such as the prefrontal cortex (PFC; mesocortical pathway), and to limbic regions, such as the amygdala (mesolimbic pathway) [11].

Neurohistological studies show that OT and DA receptor binding sites and neuronal fibers are located in close proximity [11]. The paraventricular nucleus (PVN) and supraoptic nucleus (SON) of the hypothalamus, primary OT production sites, are innervated by DA neurons and express DA receptors [12]. Conversely, DA-expressing brain regions in the mesocorticolimbic pathway receive innervation from PVN OT neurons and express both DA and OT receptors [12]. Functionally, PVN OT neurons have been shown to enhance DA activity in the VTA and suppress DA activity in the substantia nigra pars compacta (SNc) [13], an interaction that appears important for OT’s modulatory effects on social behavior [14].

In this study, we investigate the putative interdependence between OT and DA in social safety learning using the social transmission of food preference (STFP) paradigm. STFP is based on the observation that rodents (observers) display neophobia toward novel food, but develop a preference for it when a conspecific demonstrator signals its safety (i.e., social safety learning). In previous work, we proposed that STFP acquisition may involve cognitive processes that parallel functional aspects of human trust-based learning [15] and found that OT administered to male mice prior to the observer-demonstrator interaction enhanced learning in a trust acquisition condition, but blocked learning in a trust violation condition [16]. That is, mice continued to prefer demonstrated food after it was paired with lithium chloride (LiCl)-induced nausea [16].

Mesocorticolimbic DA has recently been conceptualized as a teaching signal that filters out irrelevant information by coding salience and updating expectations based on relevant inputs [9,17]. Accordingly, DA may potentiate the shift from fear to safety during social safety learning, but also respond to unexpected threats during threat learning [9,18]. Within the context of STFP acquisition, OT may modulate DA released during social interaction to enhance the salience of social information on safety [19], thereby enhancing learning in the trust acquisition condition and blocking learning in the trust violation condition. Complementary, OT may block learning in the trust violation condition by attenuating DA released in response to unexpected threats (i.e., LiCl-induced nausea).

Therefore, the current study investigated the effects of OT stimulation and DA depletion using tetrabenazine (TBZ) on social safety learning during trust acquisition (STFP1) and trust violation (STFP2) conditions. Building on our previous findings [16], we hypothesized that OT would enhance learning in the trust acquisition condition but only when DA signaling is intact. Conversely, we expected both OT stimulation and DA depletion to block learning from trust violation, OT by enhancing salience and attenuating PE processing, and DA depletion by impairing DA-mediated PE signaling. Additionally, we tested whether any effects of OT and DA system manipulations extended to other behavioral domains such as explorative and anxiety-like behavior, sociability, and spatial working memory.

## Materials and methods

### Animals

The current study included 72 male C57BL/6J mice (Janvier Labs, France), aged 10-12 weeks upon arrival, randomly divided into 3 experimental groups (OT, TBZ and OT+TBZ). Instead of testing a new vehicle control (VEH) group, we reused previously collected control data [16], a method proposed as valid to reduce the number of animals in research [20]. Mice in the VEH group (*n* = 36, unless otherwise noted) received phosphate-buffered saline (PBS) or 1% dimethyl sulfoxide (DMSO) diluted in saline (SAL) and were tested under the same conditions (same experimenters and lab environment with similar housing) as mice in the current study. Mice were housed in groups of 4 in Macrolon cages with wood-shaving bedding and cage enrichment (e.g., nesting material and toilet paper rolls to hide), kept under standard conditions (22-25 °C, humidity 50-70%, 12h light/dark cycle with lights on at 07:00 am), with ad libitum access to food and water, unless otherwise noted.

### Treatment preparation and administration

All solutions were prepared as stocks, and stored in small aliquots at -20°C. Aliquots were coded to allow blinded experiments. OT or its VEH (PBS) and TBZ or its VEH (DMSO), were administered 30 minutes and 2 hours before each behavioral test, respectively. TBZ is a selective and reversible inhibitor of vesicular monoamine transporter-2 (VMAT-2) and prevents monoamine storage with most pronounced effects on (striatal) DA [21–22]. To ensure that DA levels were replenished between behavioral tests and to minimize the risk of chronic TBZ effects, tests were scheduled at least 1 week apart.

OT was dissolved in PBS (1 µg/µL) and OT or PBS was administered intranasally (2 x 6 µL) using a micropipette [23]. To ensure proper inhalation, we applied droplets of the respective solutions gradually over the mice’s nostrils before continuing with subsequent portions. TBZ was dissolved in DMSO (7.5mg/mL in DMSO). This stock solution was diluted 100x with saline (SAL), VEH was 1% DMSO in SAL. TBZ (0.75 mg/kg) [21] or VEH was administered intraperitoneally (i.p.; 1% body weight). LiCl (1 mEq/kg) was dissolved in SAL and administered i.p. (1% body weight) 30 min after STFP2. We previously showed that this low-dose LiCl protocol effectively induced taste aversion learning [16].

### Behavioral testing

Mouse behavior was evaluated in the following order: explorative and anxiety-like behavior (open field, OF), sociability (social preference, SP), spatial working memory (T-maze), and trust-like behavior (STFP1 and STFP2). Procedures and results of control measures (OF, SP and T-maze) are reported in Supplementary file 1. Thirty minutes prior to each test, mice were habituated to the testing room. In social tests, we evaluated behavior of experimental mice towards unfamiliar mice of the same sex, strain (i.e., strangers), and age in STFP1 and STFP2 (i.e., demonstrators). To prevent transfer of unfamiliar animal scents, tested experimental mice were kept separate from naïve cage mates. For tasks involving different food types (STFP1, STFP2), gloves were changed to avoid contamination. Setups were thoroughly cleaned with 70% ethanol in between mice. To ensure blinding, treatment administration and behavioral testing were performed by different experimenters. All procedures were conducted during the light phase of mice’s cycle and approved by the Animal Ethics Committee of the University of Leuven, in accordance with EU directive 2010/63/EU on animal experiments.

### Social safety learning

We modified our STFP protocol to model trust acquisition and trust violation [24], as outlined in Budniok et al. [16]. Briefly, testing for trust acquisition (**STFP1**) and trust violation (**STFP2**) conditions was conducted in a three-compartment setup across 3 phases (*habituation*, *social interaction* and *test*) over 5 days. Throughout, experimental mice (observers) were placed on a scheduled feeding regimen (1h/day), while demonstrators were fed regular crushed food pellets mixed with paprika (STFP1) or rosemary powder (STFP2; 1% w/w) for at least 5 days before the *social interaction* phase.

Following two days of scheduled feeding, observers were habituated to the setup during a 20-min *habituation* phase on day 3. On day 4, after receiving treatment, observers interacted for 30 minutes with demonstrators placed in a round wire cage (Ø = 10 cm) in the central compartment of the setup during the *social interaction* phase (STFP1). To model trust violation (STFP2), we administered LiCl to observers 30 min after this phase to induce nausea. LiCl-induced nausea replaces the positive association with the demonstrated flavor (i.e., food is safe) with an unexpected negative association, inducing PE, which should result in decreased preference for demonstrated food. On day 5, in the *test* phase, observers had 2 h access to demonstrated (i.e., paprika in STFP1 and rosemary in STFP2) and novel food (i.e., oregano in STFP1 and basil in STFP2). Different demonstrators were used in STFP1 and STFP2, and to prevent contamination, demonstrated and novel food flavors were not counterbalanced.

Movements of observers were tracked using the ANY-Maze Video Tracking System (Stoelting, Dublin, Ireland) to assess time spent in each compartment. Additionally, we weighed food cups before and after the *test* phase to measure consumption. Social investigation was assessed as time spent near the demonstrator in the *social interaction* phase. Trust-like behavior was evaluated by comparing consumption of demonstrated versus novel food, and by calculating a demonstrated food preference score as the percentage of demonstrated food eaten (demonstrated food preference_food_ = demonstrated food eaten/(demonstrated food + novel food eaten)*100) and as time spent near demonstrated food over the whole test period (demonstrated food preference_time_ = time spent near demonstrated food/(time spent near demonstrated + novel food)*100). Demonstrated food preference_time_ was also calculated per 10-min time bin to compute the area under the curve relative to the increase from chance level (AUCi_time_) [25], allowing us to compare groups on demonstrated food preference_time_ during the first hour of the *test* phase. Higher AUCi_time_ values indicate stronger preference to spend time near demonstrated food.

### Statistical analysis

Experimental groups (OT, TBZ, OT+TBZ) were compared both with each other and individually against the VEH group (control-based comparisons). Control-based comparisons assessed mean differences (MD) using Dunnett’s *t*-test. Time spent near the demonstrator during the *social interaction* phase of STFP1 and STFP2 was not compared to the VEH group, as they had interacted with different demonstrators than the experimental groups. Comparisons among experimental groups used Welch’s *t*-tests. Within-group demonstrated food preference was evaluated by comparing demonstrated and novel food consumption within each group using paired *t*-tests, and by comparing demonstrated food preference score_time_ to chance (50%) using one-sample *t*-tests. Scores above or below chance were interpreted as increased or decreased preference, respectively.

Furthermore, we conducted exploratory analyses to assess the relative influence of observer-demonstrator interaction time in each group on preference for demonstrated food. We analyzed AUCi_time_ using two-way ANOVA, with time (below or above median: short vs. long) and group (and their interaction) as between-subjects variables. Food consumption was analyzed with three-way mixed ANOVA with the addition of food type (demonstrated vs. novel) as within-subjects variable. Interactions between all variables were also analyzed with an exploratory aim. ANOVA results were followed up with Welch’s *t*-tests.

The Benjamini-Hochberg method was applied in confirmatory analyses to control for multiple testing when assessing social safety learning (preference for demonstrated food operationalized in terms of time and consumption), both in between- and within-group comparisons [26–27]. Effect sizes were estimated using generalized eta squared (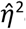G) for ANOVA effects, and Cohen’s *d* for Dunnett’s *t*-tests, Welch’s *t*-tests, paired *t*-tests, and one-sample *t*-tests. Statistical significance was set at α = 0.05, and the sample size was chosen to achieve a power of 0.80.

Data is visualized using line plots with error bars representing the standard error of the mean, and boxplot diagrams. The boxplots show the median (horizontal line in box), mean (+) and interquartile range (IQR) including the 25th percentile (Q1; lower end of box) and 75th percentile (Q3; upper end of box). Outliers (individual data points) are shown when values are higher or lower than respectively Q3 + 1.5 x IQR and Q1 – 1.5 x IQR. Extreme outliers (Q1 – 3 x IQR or Q3 + 3 x IQR) were winsorized to avoid excessive impact on analyses. Analyses were conducted in R [28].

## Results

Results of control measures (OF, SP, and T-maze) indicated that explorative and anxiety-like behavior, sociability and spatial working memory were similar between control and experimental groups, and across experimental groups (Supplementary file 1).

### Dopamine depletion reduces the enhancing effect of oxytocin on social safety learning during trust acquisition

Before starting the trust acquisition condition, one mouse from the OT group was culled due to severe wounds caused by its cage mates (*N* = 71, *n*_OT_: 23). The VEH group consisted of 34 mice. In the *social interaction* phase (Fig. 1A), all experimental groups spent a similar amount of time near the demonstrator. The following day, in the *test* phase (Fig. 1B-C), all groups preferred to spend time near, and consume, demonstrated food, indicating successful social safety learning (Supplementary file 1).

**Fig. 1.**
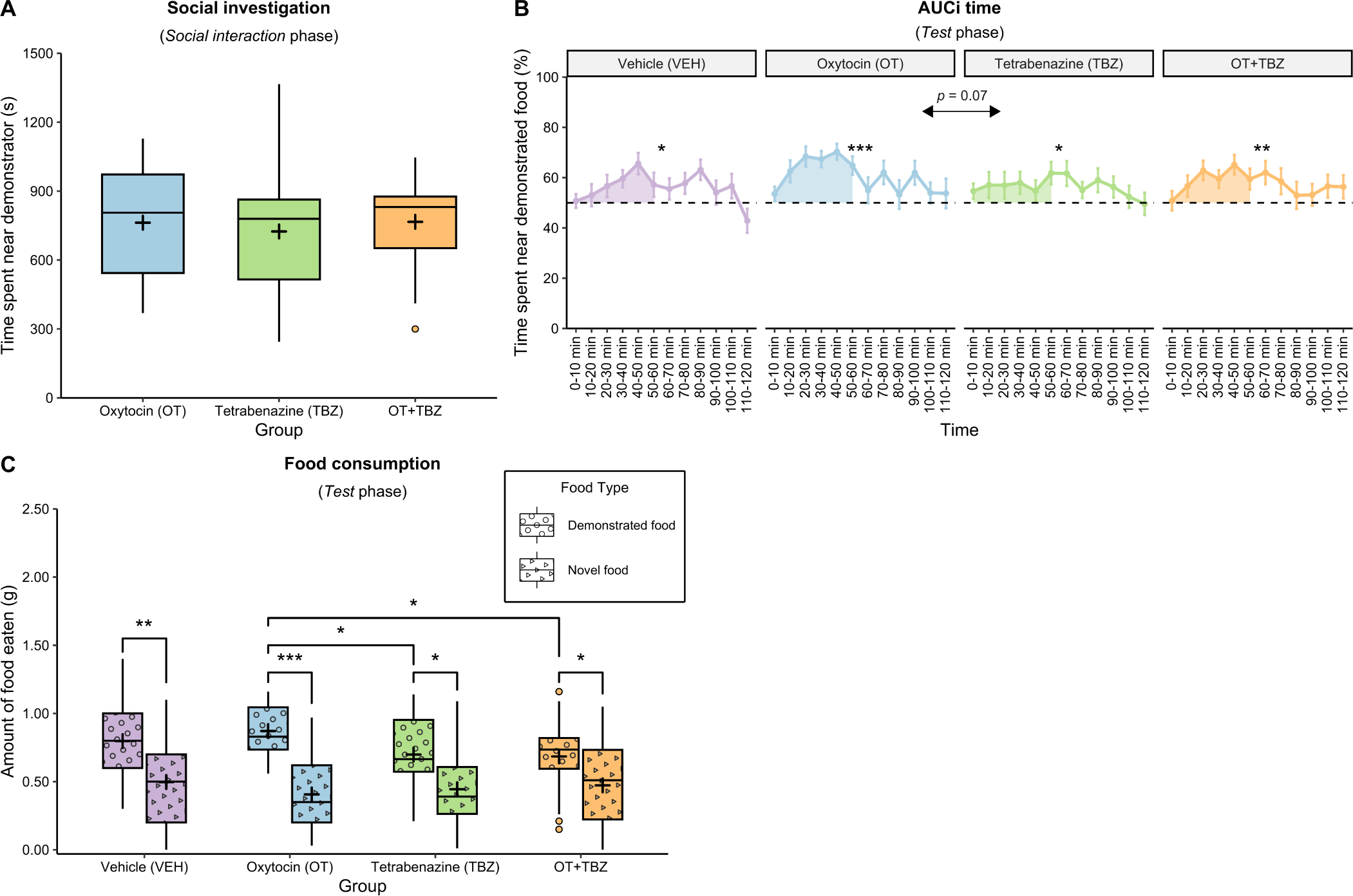
Dopamine depletion reduces the enhancing effect of oxytocin on social safety learning during trust acquisition. **(A)** In the *social interaction* phase, time spent near the demonstrator was similar for OT-treated mice and those in the TBZ (*t*_45_ = 0.49, *p* = 0.62, *d* = 0.14) and OT+TBZ groups (*t*_41_ = 0.06, *p* = 0.95, *d* = 0.02), and in the TBZ group compared to the OT+TBZ group (*t*_42.2_ = 0.62, *p* = 0.54, *d* = 0.18). **(B)** In the *test* phase, demonstrated food preference scores_time_ of all groups were significantly above chance, indicating a preference to spend time near demonstrated food (asterisks on top of line graphs). However, during the first hour of the *test* phase (shaded area), these scores tended to be higher in the OT group compared to TBZ-treated mice (*t*_39.8_ = 1.90, *p* = 0.07, *d* = 0.55). Error bars represent the standard error of the mean. AUCi = the area under the curve relative to the increase from chance level. **(C)** Similarly, all groups preferred to consume demonstrated food during the *test* phase, though this preference was enhanced for OT-treated mice compared to those in the TBZ (*t*_40_ = 2.57, *p* = 0.03, *d* = 0.75) and OT+TBZ groups (*t*_41.3_ = 2.89, *p* = 0.01, *d* = 0.84). These differences could not be attributed to enhanced total food consumption since novel food consumption was similar in the OT and TBZ groups (*t*_44.8_ = 0.49, *p* = 0.62, *d* = 0.14), and in the OT and OT+TBZ groups (*t*_44.8_ = 0.85, *p* = 0.40, *d* = 0.25). \**p* < 0.05, **p < 0.01, \*\*\**p* < 0.001

Demonstrated food preference_time_ appeared to be enhanced in the OT group during the first hour of the *test* phase (Fig. 1B), so we calculated AUCi_time_ over the first hour of the *test* phase to compare groups. Control-based comparisons revealed no differences in preference to spend time near, and consume, demonstrated food between the VEH group and any of the experimental groups, indicating similar social safety learning. In contrast, AUCi_time_ of the OT group tended to be higher compared to the TBZ group. Consumption of demonstrated food was significantly increased in the OT group, both compared to mice treated with TBZ, and OT+TBZ. These differences could not be attributed to enhanced total food consumption since novel food consumption was similar in all experimental groups.

### Oxytocin enhances social safety learning during trust acquisition specifically in mice with shorter interaction times

OT enhanced social safety learning compared to experimental groups, but not compared to previously collected control data. STFP acquisition is influenced by the amount of observer exposure to a specific demonstrator, with observers that have shorter interaction times being less likely to develop a preference for demonstrated food [29]. OT’s effects may be more pronounced in observers with shorter interaction times. Therefore, a post-hoc analysis categorized observers into groups with longer (x > median) and shorter (x ≤ median) interaction times by calculating the median interaction time over all experimental groups but separately within the previous (median = 911.7s) [16] and current experiments (median = 807.3s). Since we did not conduct such an analysis on the OT group of our previous experiment (OT_prev_) [16], we included this group in the analysis as well (*n* = 12). Splitting all groups based on the median resulted in approximately equal-sized subgroups (see Table 1).

**Table 1.** Distribution of mice within each group after median split based on interaction time with the demonstrator.

| Condition | Group | Interaction time |  |
| --- | --- | --- | --- |
|  |  | *Short* | *Long* |
| Trust acquisition (STFP1) | Vehicle (VEH) | 15 | 19 |
|  | Oxytocin previous experiment (OT<sub>prev</sub>) | 6 | 6 |
|  | Oxytocin (OT) | 12 | 11 |
|  | Tetrabenazine (TBZ) | 13 | 11 |
|  | OT + TBZ | 11 | 13 |
| Trust violation (STFP2) | Vehicle (VEH) | 18 | 16 |
|  | Oxytocin previous experiment (OT<sub>prev</sub>) | 5 | 6 |
|  | Oxytocin (OT) | 11 | 12 |
|  | Tetrabenazine (TBZ) | 13 | 11 |
|  | OT + TBZ | 11 | 11 |

*Note.* The median interaction time was calculated over all experimental groups but separately within the previous (VEH and OT<sub>prev</sub>) [16] and current experiments (OT, TBZ, OT+TBZ).

A two-way ANOVA on AUCi_time_ (Fig. 2A) revealed a significant interaction between group and interaction time (*F*_4,107_ = 3.16, *p* = 0.02, 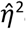G = 0.11). When interaction time was longer, we found no differences between the VEH group and any experimental group, nor across experimental groups. When interaction time was shorter, only OT-treated groups had a demonstrated food preference_time_ above chance independent of DA availability. Moreover, AUCi_time_ of mice in both the OT and OT_prev_ groups was significantly higher compared to mice in the VEH and TBZ groups. OT_prev_ also increased AUCi_time_ compared to the OT+TBZ group, while AUCi_time_ in the OT+TBZ group tended to be higher than in the TBZ group.

**Fig. 2.**
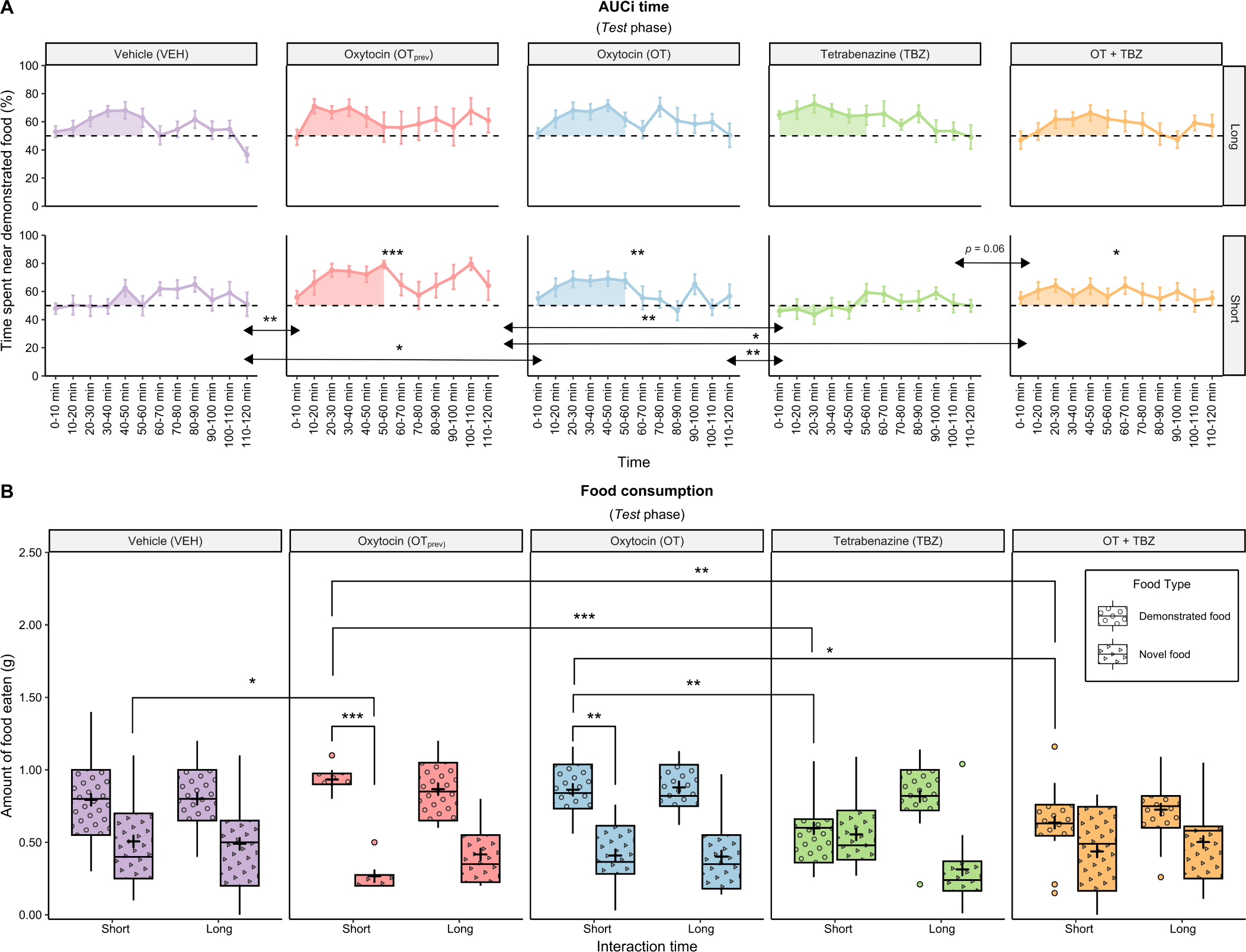
Oxytocin enhances social safety learning during trust acquisition specifically in mice with shorter interaction times. **(A)** Exploratory analyses revealed in mice with longer interaction times no differences in AUCi_time_ (shaded area) between the VEH group and any experimental group, nor across the experimental groups. When interaction time was shorter, only OT-treated groups had a demonstrated food preference_time_ above chance independent of DA availability (OT: *t*_11_ = 3.14, *p* = 0.009, *d* = 0.91; OT_prev_: *t*_5_ = 10.1, *p* < 0.001, *d* = 4.12; OT+TBZ: *t*_10_ = 2.58, *p* = 0.03, *d* = 0.78; asterisks on top of line graphs). Moreover, AUCi_time_ of mice in both the OT and OT_prev_ group was significantly higher compared to mice in the VEH (OT vs VEH: *t*_24.8_ = 2.31, *p* = 0.03, *d* = 0.88; OT_prev_ vs VEH: *t*_18.5_ = 3.32, *p* = 0.004, *d* = 1.36) and TBZ groups (OT vs TBZ: *t*_22_ = 2.93, *p* = 0.008, *d* = 1.17; OT_prev_ vs TBZ: *t*_16.9_ = 3.94, *p* = 0.001, *d* = 1.70). OT_prev_ also increased AUCi_time_ compared to the OT+TBZ group (*t*_14.4_ = 2.14, *p* = 0.049, *d* = 1.02), while AUCi_time_ in the OT+TBZ group tended to be higher than in the TBZ group (*t*_21.6_ = 1.99, *p* = 0.06, *d* = 0.81). Error bars represent the standard error of the mean. AUCi = the area under the curve relative to the increase from chance level. **(B)** When interaction time was longer, we found no differences in consumption of demonstrated food between the VEH group and any experimental group, nor across experimental groups. When interaction time was shorter, only mice in the OT (*t*_11_ = 4.11, *p* = 0.002, *d* = 1.19) and OT_prev_ (*t*_5_ = 9.33, *p* < 0.001, *d* = 3.81) groups had a preference to consume demonstrated food. This preference was enhanced in the OT_prev_ group compared to VEH-treated mice, as consumption of demonstrated food tended to be higher and novel food consumption significantly lower (*t*_18.98_ = -2.55, *p* = 0.02, *d* = 1.02). Differences between the OT and VEH groups trended in the same direction, but did not reach statistical significance. Lastly, mice in both the OT and OT_prev_ groups consumed significantly more demonstrated food than mice in the TBZ (OT vs TBZ: *t*_22.2_ = 3.02, *p* = 0.006, *d* = 1.20; OT_prev_ vs TBZ: *t*_17_ = 4.16, *p* < 0.001, *d* = 1.76) and OT+TBZ groups (OT vs OT+TBZ: *t*_17_ = 2.21, *p* = 0.04, *d* = 0.93; OT_prev_ vs OT+TBZ: *t*_13.8_ = 3.08, *p* = 0.008, *d* = 1.38). \**p* < 0.05, \*\**p* < 0.01, \*\*\**p* < 0.001

A three-way mixed ANOVA on food consumption (Fig. 2B) with the addition of food type (demonstrated vs. novel) as within-subjects variable revealed no significant interaction between group, interaction time and food type (*F*_4,107_ = 1.30, *p* = 0.27, 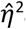G = 0.04). However, a visual inspection of the data revealed a pattern similar to the results for AUCi_time_ (Fig. 2A). When interaction time was longer, we found no differences between the VEH group and any experimental group, nor across experimental groups. When interaction time was shorter, only mice in the OT, and OT_prev_ groups had a preference to consume demonstrated food. This preference was enhanced in the OT_prev_ group compared to VEH-treated mice, as consumption of demonstrated food tended to be higher and novel food consumption significantly lower. Differences between the OT and VEH groups trended in the same direction, but did not reach statistical significance. Mice in both the OT and OT_prev_ groups consumed significantly more demonstrated food than mice in the TBZ, and OT+TBZ groups. Together, these findings indicated that OT enhanced social safety learning specifically in mice that interacted shorter with the demonstrator.

### Oxytocin requires intact dopaminergic signaling to block learning during trust violation

Before starting the trust violation condition, two mice from the OT+TBZ group were culled due to severe wounds caused by their cage mates (*N* = 69, *n*_OT+TBZ_: 22). The VEH group consisted of 34 mice. In the *social interaction* phase (Fig. 3A), all experimental groups spent a similar amount of time near the demonstrator. The following day, in the *test* phase (Fig. 3B-C), only mice in the OT group preferred to spend time near, and consume, demonstrated food. In contrast, mice in the VEH and OT+TBZ groups had no such preference, while TBZ-treated mice preferred to consume demonstrated food but not to spend time near it (Supplementary file 1).

**Fig. 3.**
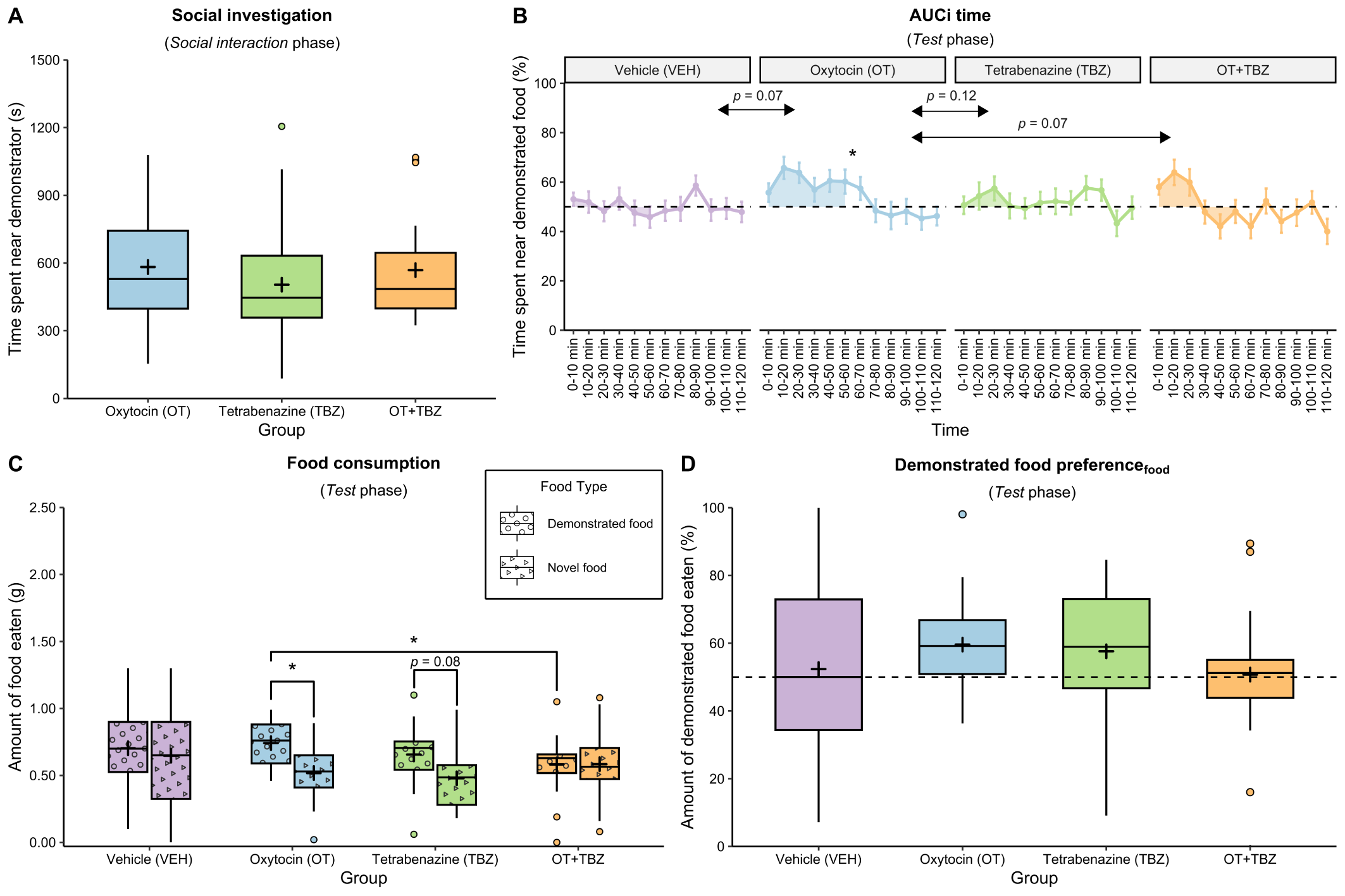
Oxytocin requires intact dopaminergic signaling to block learning during trust violation. **(A)** In the *social interaction* phase, time spent near the demonstrator before experiencing LiCl-induced nausea was similar for OT-treated mice and those in the TBZ (*t*_44.9_ = 1.03, *p* = 0.31, *d* = 0.30) and OT+TBZ (*t*_42.9_ = 0.19, *p* = 0.85, *d* = 0.06) groups, and in the TBZ group compared to the OT+TBZ group (*t*_44_ = 0.89, *p* = 0.38, *d* = 0.26). **(B)** In the *test* phase, only mice in the OT group had a demonstrated food preference score_time_ significantly above chance, indicating a preference to spend time near demonstrated food (asterisk on top of line graph). This preference tended to be higher during the first hour of the *test* phase (shaded area) compared to the VEH (MD = 54.56, *p* = 0.07, *d* = 0.71), OT+TBZ (*t*_41.1_ = 1.84, *p* = 0.07, *d* = 0.55) and TBZ groups (*p* = 0.06), but the latter effect did not survive multiple testing correction (*t*_42.4_ = 1.94, *p* = 0.12, *d* = 0.56). Error bars represent the standard error of the mean. AUCi = the area under the curve relative to the increase from chance level. **(C)** Only mice in the OT and TBZ groups retained preference to consume demonstrated food, and this preference was enhanced for OT-treated mice compared to those in the OT+TBZ group (*t*_41.8_ = 2.76, *p* = 0.02, *d* = 0.83). This difference could not be attributed to enhanced total food consumption since novel food consumption was similar in the OT and OT+TBZ groups (*t*_41.8_ = 0.95, *p* = 0.35, *d* = 0.28). **(D)** Control-based comparisons failed to indicate differences in demonstrated food preference_food_ between the VEH group and any of the experimental groups. \**p* < 0.05

The loss of demonstrated food preference in the VEH group appeared to be driven by an increase in novel food consumption compared to the experimental groups (Fig. 3C). Therefore, control-based comparisons used AUCi_time_ (Fig. 3B) and demonstrated food preference_food_ (Fig. 3D) instead of demonstrated food consumption. AUCi_time_ tended to be higher in OT-treated mice compared to mice in the VEH group but preference to consume demonstrated food was similar.

Among experimental groups, AUCi_time_ tended to be higher in the OT group compared to mice in the OT+TBZ and TBZ groups. Consumption of demonstrated food was similar for OT-treated mice compared to mice in the TBZ group, but enhanced compared to mice in the OT+TBZ group. This difference could not be attributed to enhanced total food consumption since novel food consumption was similar in the OT and OT+TBZ groups.

### Trust violation reduces preference for demonstrated food specifically in mice with longer interaction times

In the trust violation condition, preference to spend time near, and consume, demonstrated food was reduced in the VEH, TBZ (only in terms of time) and OT+TBZ groups. Given the effect of interaction time on preference for demonstrated food in the trust acquisition condition, we conducted a post hoc analysis to explore whether trust violation specifically influenced preference for demonstrated food in mice with shorter or longer interaction times with the demonstrator.

Similar as in the trust acquisition condition, we split the treatment groups based on the median of our previous (median = 884.6s) [16] and current experiments (median = 498.9s), and included the OT group of our previous experiment (OT_prev_) [16] as well (*n* = 11). This resulted in approximately equal-sized subgroups (see Table 1).

A two-way ANOVA on AUCi_time_ (Fig. 4A) revealed no significant interaction between group and interaction time (*F*_4,104_ = 0.54, *p* = 0.71, 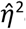G = 0.02). Similarly, a three-way ANOVA on food consumption (Fig. 4B) revealed no significant interaction between group, interaction time and food type (*F*_4,104_ = 0.33, *p* = 0.86, 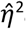G = 0.01). In line with the above-reported effects of the current OT group, OT_prev_ retained preference to spend time near, and consume, demonstrated food. These results indicated that mice with longer interaction times in the VEH, TBZ, and OT+TBZ groups, who had developed a preference for demonstrated food in the trust acquisition condition, lost this preference when we administered LiCl (i.e., trust violation) after the observer-demonstrator interaction. This attenuated the preference to spend time near, and consume, demonstrated food at the group level in the VEH, TBZ (only in terms of time), and OT+TBZ groups, while mice receiving only OT were less affected by LiCl-induced nausea and retained preference for demonstrated food.

**Fig. 4.**
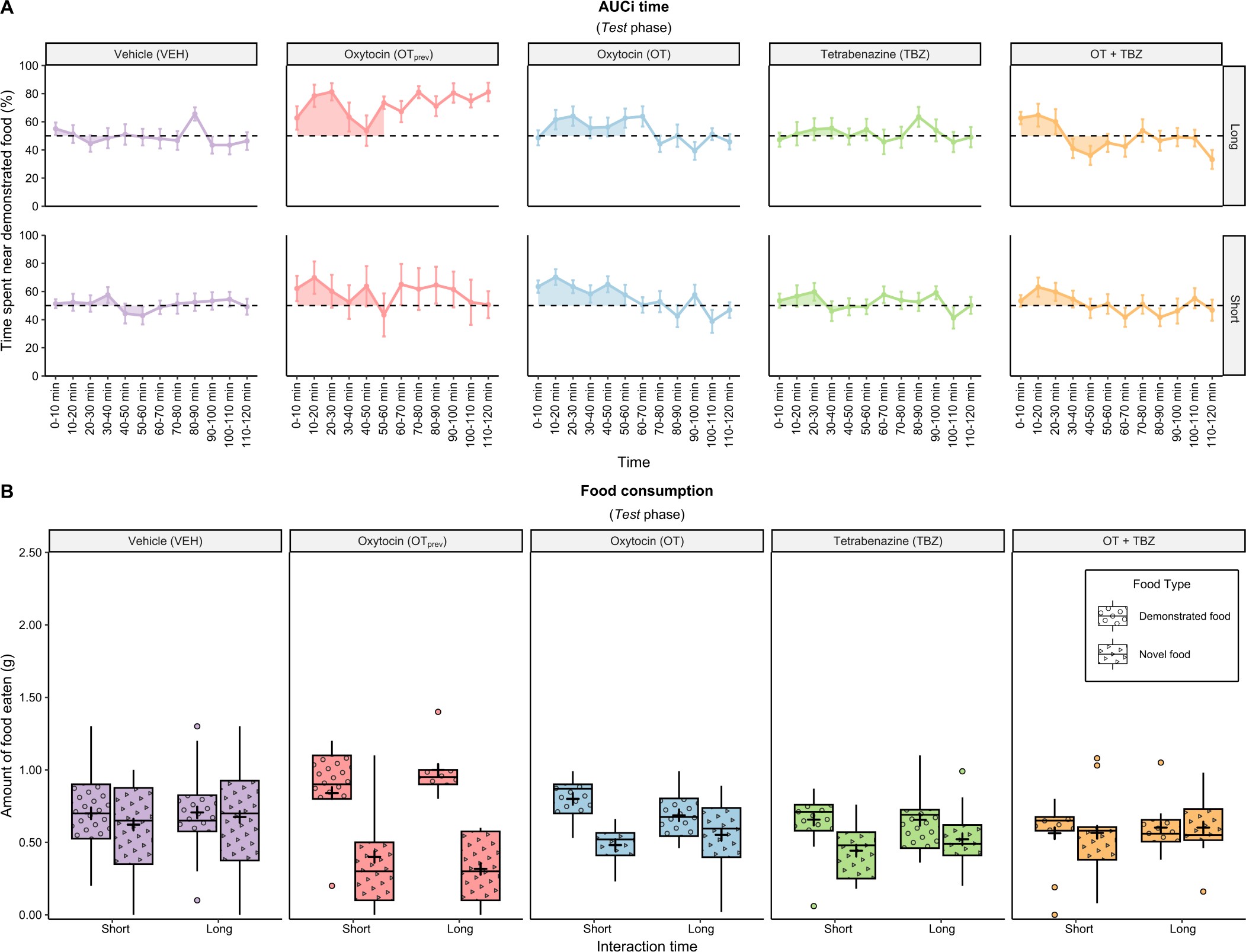
Trust violation reduces preference for demonstrated food specifically in mice with longer interaction times. Exploratory analyses revealed that mice with longer interaction times with the demonstrator in the VEH, TBZ and OT+TBZ groups, who had developed a preference for demonstrated food in the trust acquisition condition, lost this preference when we administered LiCl (i.e., trust violation) after the observer-demonstrator interaction. This attenuated the preference to **(A)** spend time near, and **(B)** consume, demonstrated food at the group level in the VEH, TBZ (only in terms of time), and OT+TBZ groups, while mice receiving only OT, in our current (see Figure 3) and previous experiments (time: *t*_10_ = 2.91, *p* = 0.02, *d* = 0.88; food: *t*_10_ = 3.04, *p* = 0.02, *d* = 0.92), were less affected by LiCl-induced nausea and retained preference for demonstrated food. Error bars represent the standard error of the mean. AUCi = the area under the curve relative to the increase from chance level.

## Discussion

We investigated whether DA is required for OT to modulate social safety learning using the STFP paradigm, interpreting STFP acquisition as a putative functional parallel to human trust-based learning [15–16]. As hypothesized, OT enhanced learning in the trust acquisition condition only when DA signaling was intact. In the trust violation condition, OT stimulation and DA depletion blocked learning, with retained preference for demonstrated food as result, except when OT was administered under DA depletion. Exploratory analyses in the trust acquisition condition revealed that OT enhanced preference for demonstrated food specifically in mice with shorter interaction times, a group that failed to develop this preference when not treated with OT, or when treated with OT under DA depletion. In the trust violation condition, the loss of preference for demonstrated food in the VEH, TBZ, and OT+TBZ groups was driven by reduced preference in mice with longer interaction times.

Although OT produced similar behavioral outcomes across OT groups, differences between VEH and the current OT group did not reach statistical significance, unlike in our previous experiment [16]. STFP acquisition depends on associating a food odor with carbon disulfide (CS_2_), a semiochemical in the demonstrator’s breath that signals safety [30]. Our results suggest that OT enhanced the salience of this association via DA modulation, particularly in mice for whom this association was less salient [5–6]. This modulation may occur in the amygdala, a key node in salience processing that integrates socially-relevant sensory information and expresses both OT and DA receptors [31–33]. The discrepancy in strength of OT’s salience effect between our experiments may stem from different demonstrators used, which is consistent with recent findings showing that OT’s effects on STFP acquisition are modulated by demonstrator characteristics [34].

We assessed social safety learning through preference for demonstrated food, measured by time spent near, and consumption of, demonstrated food. In the trust acquisition condition, OT increased time spent near demonstrated food, independent of DA depletion (Fig. 2A), but enhanced consumption only when DA was available (Fig. 1C). The incentive-sensitization theory [35–36] may explain this pattern of results, as it distinguishes between ‘liking’ (value, DA-independent) and ‘wanting’ (motivational drive, DA-dependent) of a reward like demonstrated food. OT mediates entrainment of neutral odors to appetitive social cues [37], suggesting that under DA depletion, OT may still have modulated ‘liking’ of demonstrated food odor by increasing its social relevance (i.e., neutral food odor becomes social odor). In contrast, without DA, OT may have failed to enhance the association between food odor and safety (CS_2_ signal) required to increase ‘wanting’ (or consumption) of demonstrated food (i.e., social safety learning).

In the trust violation condition, OT retained preference to spend time near (‘liking’; Fig. 3B) and consume demonstrated food (‘wanting’; Fig. 3C), but only when DA was available, while DA depletion alone retained preference to consume demonstrated food (‘wanting’) but not spend time near it (‘liking’; Fig. 3C). Under DA depletion, ‘liking’ may have been reduced, as mice tend to avoid neutral and social odors paired with LiCl [38–39], while ‘wanting’ persisted due to impaired PE updating [9]. Although OT enhances avoidance of negative stimuli to reduce ‘liking’ (e.g., demonstrated food odor paired with LiCl; [37]), this effect may have been blocked as OT first supported the positive association between the food odor and safety (CS_2_ signal) and complementary attenuated PE processing (LiCl-induced nausea), to increase ‘wanting’, mitigating LiCl effects, but only when DA was available.

Since salience processing is not exclusively attributable to DA [40–41], our results suggest that OT modulates ‘liking’ and ‘wanting’ of social stimuli via distinct neural mechanisms. This differential modulation could have important implications for therapeutic interventions. For instance, OT has been proposed as adjunct pharmacotherapy to treat psychiatric disorders characterized by social deficits such as autism spectrum disorder (ASD; e.g., [42–43]). However, the success of OT in treatment of ASD has been variable [44], which may reflect the heterogeneity of ASD etiology and the existence of different ASD subtypes. Our findings suggest that OT’s efficacy may be reduced in ASD subtypes involving DA deficits [45].

In conclusion, our results confirm the involvement of OT in social safety learning, demonstrating that OT modulates DA signaling to enhance trust acquisition and block learning from trust violation by complementary enhancing the salience of social information on safety and reducing PE processing. Notably, OT may exert its salience effect particularly in mice for whom social information is less salient, though further research is required to confirm this hypothesis.

Furthermore, the incentive-sensitization theory [35–36] provides a valuable framework for distinguishing between aspects of social behavior OT influences via DA and those it influences through other neurochemical systems. Given OT’s therapeutic potential (e.g., [42,46]), elucidating the neurochemical pathways underlying its effects is crucial to optimize clinical outcomes.

## Author contributions

Samuel Budniok: Investigation, Data Curation, Visualization, Formal analysis, Writing - Original Draft,

Writing - Review & Editing.

Zsuzsanna Callaerts-Vegh: Conceptualization, Formal analysis, Writing - Review & Editing, Supervision.

Marian Bakermans-Kranenburg: Conceptualization, Formal analysis, Writing - Review & Editing.

Guy Bosmans: Conceptualization, Formal analysis, Writing - Review & Editing, Funding acquisition.

Rudi D’Hooge: Conceptualization, Formal analysis, Writing - Review & Editing, Funding acquisition, Supervision.

