## Supplementary file 1 for "Interdependency between oxytocin and dopamine in trust-based learning in mice"

**Methods**

**Behavioral testing**

***Tests of explorative and anxiety-like behavior, and spatial working memory***

**Open field (OF).** Explorative and anxiety-like behavior were evaluated by introducing dark-acclimated mice into a brightly lit Plexiglass arena (50 x 50 cm) placed inside a cupboard for 10 minutes. During the test, mouse movements were recorded by an overhead camera connected to the ANY-Maze Video Tracking System (Stoelting, Dublin, Ireland). Explorative behavior was quantified by measuring total path length (in m), while anxiety-like behavior was evaluated based on time spent in the arena's center versus its periphery (thigmotaxis). A greater duration in the center is generally interpreted as a sign of reduced anxiety-like behavior in rodents [47].

**Spatial working memory T-maze.** Spatial working memory was evaluated in a T-maze. The maze consists of 3 arms made of dark grey plastic arranged in a T, with 2 goal arms (47 cm long, 6 cm wide, enclosed by 4 cm walls) and one start arm (50 cm long, 6 cm wide, enclosed by 4 cm walls), and is placed on legs to elevate the maze (40 cm above the table). The goal arms can be blocked by a barrier (10 cm within arm). Mouse movements were recorded by an overhead camera connected to a tracking software (Ethovision, Noldus, Wageningen).

The protocol consisted of 2 consecutive 10-min trials, each beginning with the placement of a mouse in the start arm. During the first trial, only one goal arm was accessible (familiar arm; the position of the familiar arm was counterbalanced between groups). Mice, as natural scavengers are inclined to explore new areas, which requires memory of locations they already visited. Therefore, we placed the mouse briefly in a holding cage between trials while cleaning the maze thoroughly with 70% ethanol and removing the barrier. The mouse was then returned to the start arm for the second trial. Driven by their scavenging instinct, mice typically prefer exploring the newly accessible arm (novel arm). Path length (in m) in both trials was recorded to evaluate activity levels, and we calculated a novel arm preference score during trial 2 (time spent in novel arm/(time spent in novel + familiar arm)*100). This score evaluated mice’s ability to flexibly adapt their behavior based on information in the working memory.

***Test of sociability***

**Social preference (SP).** Sociability was evaluated in a three-compartment setup. The setup consists of a central chamber (40 cm long, 10 cm wide), and 2 side chambers (10 x 10 cm) made from transparent Plexiglass. The chambers were separated by perforated walls (1 cm diameter holes), allowing limited tactile, olfactory, visual and auditory contact. Movements of the experimental mouse were recorded by an overhead camera connected to the ANY-Maze Video Tracking System (Stoelting, Dublin, Ireland).

The protocol consisted of 2 consecutive episodes, each beginning with the placement of an experimental mouse in the central chamber of the setup. Following a 5-min habituation period (Episode 1), a stranger mouse (S1) was introduced into one of the side chambers to record approach of the experimental mouse to S1 over the next 10 minutes (Episode 2). The position of S1 was counterbalanced between groups. After Episode 2, the experimental and stranger mouse were returned to their respective home cages, and the setup was thoroughly cleaned with 70% ethanol. We recorded path length (in m) per episode to evaluate activity levels and calculated an S1 preference score in terms of time spent in the different zones during Episode 2 (time spent near S1/(time spent near S1 + near empty zone)*100) to obtain a measure of sociability.

**Statistical analysis**

Activity levels during the OF test were compared between experimental groups (OT, TBZ, OT+TBZ) with Welch’s *t*-tests, while activity during the T-maze and SP tests was analyzed using two-way mixed ANOVA, with experimental group as between-subjects variable and trial (T-maze; 1 vs. 2) or episode (SP; 1 vs. 2) as within-subjects variables. Interactions between variables were also analyzed. Since activity levels may be more influenced by contextual factors, we excluded the vehicle control (VEH) group from these analyses.

For all other data, control-based comparisons used Dunnett’s *t*-test to assess mean differences (MD) between the VEH group and each experimental group (OT, TBZ, OT+TBZ), while comparisons between experimental groups were performed using Welch’s *t*-tests. In addition, the S1 (SP test) and novel arm preference (T-Maze test) scores of all groups were compared to chance (50%) using one-sample *t*-tests. Scores significantly above or below chance were interpreted as indicating a significantly increased or decreased preference, respectively. The Benjamini-Hochberg method was used to correct for multiple testing across measures assessing anxiety-like behavior (time spent in the center and periphery in the OF test) [26-27]. Generalized eta squared ($\hat{\eta}$^2^_G_) was used as an estimate of the effect size for ANOVA effects, and Cohen’s *d* for one-sample, Welch’s and Dunnett’s *t*-tests. Statistical significance was set at α = 0.05.

**Results**

***Social safety learning***

**STFP1 (trust acquisition condition).** Within-group analyses in the *test* phase revealed that all groups preferred to spend time near (VEH: *t*_33_ = 2.32, *p* = 0.03, *d* = 0.40; OT: *t*_22_ = 4.82, *p* < 0.001, *d* = 1.01; TBZ: *t*_23_ = 2.33, *p* = 0.03, *d* = 0.48; OT+TBZ: *t*_23_ = 3.18, *p* = 0.008, *d* = 0.65), and consume (VEH: *t*_33_ = 3.08; *p* = 0.008, *d* = 0.53; OT: *t*_22_ = 5.93, *p* < 0.001, *d* = 1.24; TBZ: *t*_23_ = 2.37, *p* = 0.03, *d* = 0.48; OT+TBZ: *t*_23_ = 2.19, *p* = 0.04, *d* = 0.45), demonstrated food, indicating successful social safety learning.

**STFP2 (trust violation condition).** Within-group analyses in the *test* phase revealed that only mice in the OT group preferred to spend time near (*t*_22_ = 2.23, *p* = 0.04, *d* = 0.46), and consume (*t*_22_ = 3.10, *p* = 0.01, *d* = 0.65), demonstrated food. In contrast, mice in the VEH (time: *t*_33_ = -0.01, *p* = 0.99, *d* = 0.002; food: *t*_33_ = 0.53; *p* = 0.99, *d* = 0.09), and OT+TBZ groups (time: *t*_21_ = -0.04, *p* = 0.99, *d* = 0.01; food: *t*_21_ = -0.01, *p* = 0.99, *d* = 0.003) had no such preference, while TBZ-treated mice preferred to consume demonstrated food (*t*_23_ = 2.18, *p* = 0.08, *d* = 0.45) but not spend time near it (*t*_23_ = 0.77, *p* = 0.45, *d* = 0.16).

***Tests of explorative and anxiety-like behavior, and spatial working memory***

**Open Field.** During the OF test, all experimental groups had similar pathlengths (OT vs. TBZ: *t*_42.1_ = 0.98, *p* = 0.33, *d* = 0.28; OT vs. OT+TBZ: *t*_45.8_ = 0.85, *p* = 0.40, *d* = 0.25; TBZ vs OT+TBZ: *t*_40.5_ = 0.30, *p* = 0.76, *d* = 0.09). Furthermore, time spent in the center of the arena was not significantly different between the VEH group and OT- (MD = 3.87, *p* = 0.93, *d* = 0.13), TBZ- (MD = 5.63, *p* = 0.99, *d* = 0.19), or OT+TBZ-treated mice (MD = 6.38, *p* = 1, *d* = 0.21). There were also no significant differences between mice treated with OT and TBZ (*t*_45.5_ = 1.40, *p* = 0.34, *d* = 0.40), OT and OT+TBZ (*t*_44.2_ = 0.35, *p* = 0.73, *d* = 0.10) and TBZ and OT+TBZ (*t*_45.5_ = 1.59, *p* = 0.24, *d* = 0.46).

Regarding time spent in the periphery, VEH-treated mice did not significantly differ from mice in the OT (MD = 8.98, *p* = 0.93, *d* = 0.20), TBZ (MD = 3.17, *p* = 0.99, *d* = 0.06) or OT+TBZ group (MD = 1.54, *p* = 1, *d* = 0.04). Similarly, time spent in the periphery was not significantly different for OT-treated mice compared to mice in the TBZ (*t*_42_ = 0.83, *p* = 0.41, *d* = 0.24) and OT+TBZ group (*t*_43.9_ = 0.68, *p* = 0.73, *d* = 0.20), and for TBZ- compared to OT+TBZ-treated mice (*t*_37_ = 0.34, *p* = 0.73, *d* = 0.10). These results indicate that explorative and anxiety-like behavior was similar across all groups.

**T-Maze.** A two-way mixed ANOVA on path length with experimental group (OT vs. TBZ vs. OT+TBZ) as between-subjects variable and trial (1 vs. 2) as within-subjects variable revealed neither an interaction between experimental group and trial (*F*_2,69_ = 2.92, *p* = 0.06, $\hat{\eta}$^2^_G_ = 0.02), nor a main effect of experimental group (*F*_2,69_ = 1.82, *p* = 0.17, $\hat{\eta}$^2^_G_ = 0.04). However, path length was significantly higher during trial 1 compared to trial 2 (*F*_1,69_ = 12.22, *p* < 0.001, $\hat{\eta}$^2^_G_ = 0.03) indicating that activity levels were similarly elevated across experimental groups during trial 1.

Furthermore, mice treated with VEH (*t*_23_ = 7.37, *p* < 0.001, *d* = 1.50), OT (*t*_23_ = 5.22, *p* < 0.001, *d* = 1.07), TBZ (*t*_23_ = 8.56, *p* < 0.001, *d* = 1.75) and OT+TBZ (*t*_23_ = 9.99, *p* < 0.001, *d* = 2.04) had novel arm preference scores significantly above chance, indicating that all groups preferred to explore the novel arm during trial 2. However, novel arm preference scores were not significantly different between the VEH group and mice treated with OT (MD = 4.66, *p* = 0.19, *d* = 0.50), TBZ (MD = 2.10, *p* = 0.76, *d* = 0.22) or OT+TBZ (MD = 2.05, *p* = 0.78, *d* = 0.23). Comparisons between experimental groups revealed that novel arm preference scores of the TBZ and OT+TBZ groups were similar (*t*_44.9_ = 0.02, *p* = 0.98, *d* = 0.01), but significantly increased compared to the OT group (TBZ vs. OT: *t*_46_ = 2.51, *p* = 0.02, *d* = 0.72; OT+TBZ vs. OT: *t*_45.3_ = 2.68, *p* = 0.01, *d* = 0.77). These results indicate intact spatial working memory abilities across groups, but modulation of performance by dopamine.

***Test of sociability***

**SP.** A two-way mixed ANOVA on path length with experimental group (OT vs. TBZ vs. OT+TBZ) as between-subjects variable and episode (1 vs. 2) as within-subjects variable revealed neither an interaction between experimental group and trial (*F*_2,69_ = 1.45, *p* = 0.24, $\hat{\eta}$^2^_G_ = 0.01), nor a main effect of experimental group (*F*_2,69_ = 0.08, *p* = 0.92, $\hat{\eta}$^2^_G_ = 0.002). However, path length was significantly higher during Episode 1 compared to Episode 2 (*F*_1,69_ = 150.97, *p* < 0.001, $\hat{\eta}$^2^_G_ = 0.31) indicating that activity levels similarly reduced across groups upon introduction of a social stimulus in the setup. Furthermore, mice treated with VEH (*t*_35_ = 3.55, *p* = 0.001, *d* = 0.59), OT (*t*_23_ = 2.85, *p* = 0.01, *d* = 0.58), TBZ (*t*_23_ = 3.72, *p* = 0.001, *d* = 0.76) and OT+TBZ (*t*_23_ = 2.75, *p* = 0.01, *d* = 0.56) had S1 preference scores significantly above chance, indicating that all groups preferred to spent time near the stranger. However, S1 preference scores were not significantly different between the VEH group and mice treated with OT (MD = 1.41, *p* = 0.97, *d* = 0.10), TBZ (MD = 2.72, *p* = 0.81, *d* = 0.21) or OT+TBZ (MD = 0.90, *p* = 0.99, *d* = 0.07). S1 preference scores did also not differ across experimental groups (OT vs TBZ: *t*_45.3_ = 0.32, *p* = 0.75, *d* = 0.09; OT vs OT+TBZ: *t*_46_ = 0.12, *p* = 0.91, *d* = 0.03; TBZ vs OT+TBZ: *t*_45.5_ = 0.45, *p* = 0.66, *d* = 0.13). These results indicate similar sociability across all groups.
